# Microtubule binding of the human HAUS complex is directly controlled by importins and Ran-GTP

**DOI:** 10.1101/2023.02.19.529112

**Authors:** Kseniya Ustinova, Felix Ruhnow, Maria Gili, Thomas Surrey

## Abstract

Mitotic spindle assembly during cell division is a highly regulated process. Ran-GTP produced around chromosomes controls the activity of a multitude of spindle assembly factors by releasing them from inhibitory interaction with importins. A major consequence of Ran-GTP regulation is the stimulation of local microtubule nucleation around chromosomes via augmin/HAUS-mediated branched microtubule nucleation, a process that is critically important for correct spindle assembly. However, augmin is not known to be a direct target of the Ran-GTP pathway, raising the question of how its activity is controlled. Here we present the *in vitro* reconstitution of Ran-GTP-regulated microtubule binding of the human HAUS complex. We demonstrate that importins directly bind to the HAUS complex, which prevents HAUS from binding to microtubules. Ran-GTP relieves this inhibition. Therefore, the HAUS complex is a direct target of the Ran-GTP pathway, suggesting that branching microtubule nucleation is directly regulated by the Ran-GTP gradient around chromosomes in dividing cells.

## INTRODUCTION

Mitotic spindle assembly and function rely on the precise spatiotemporal regulation of the nucleation of microtubules, their dynamic properties, and their spatial organization. While centrosomes are a major source of microtubule nucleation, during cell division microtubules are also nucleated in the vicinity of chromosomes. Errors in this nucleation pathway lead to spindle defects and chromosome segregation errors, causing cancer or aneuploidy (Cimini et al. 2001, Asteriti et al. 2010). Chromatin-mediated nucleation is controlled by Ran-GTP that is generated at the chromosomes (Oh et al. 2016). Ran-GTP displaces inhibitory importins from nuclear localization signal (NLS) containing proteins (Kalab et al. 2002, Clarke and Zhang 2008, Kalab and Heald 2008). Several critical spindle proteins, also called spindle assembly factors (SAFs), are regulated in this manner (Gruss et al. 2001, Wiese et al. 2001, Koffa et al. 2006, Ribbeck et al. 2006, Ems-McClung et al. 2004).

Microtubule nucleation around chromosomes requires the recruitment of the major microtubule nucleator γ-tubulin ring complex (γTURC) to pre-existing microtubules by the HAUS/augmin complex (Stearns and Kirschner 1994, Moritz et al. 2000, Prosser and Pelletier 2017). This nucleation mechanism leads to branched microtubule nucleation with freshly nucleated microtubules being oriented roughly parallel to preexisting ones (Petry et al. 2013). However, neither HAUS nor γTURC is known to interact with importins, and hence are not thought to be direct targets of the Ran-GTP pathway.

Instead, work mostly in *Xenopus* egg extract, has shown that branched microtubule nucleation is stimulated by the SAF TPX2 (Petry et al. 2013, Gruss et al. 2002, Alfaro-Aco et al. 2017) that has been proposed to stimulate branched microtubule nucleation by activating Aurora A kinase (Eyers and Maller 2004, Bayliss et al. 2003) which in turn may phosphorylate γTURC regulators (Pinyol et al. 2013) or by binding directly to augmin/HAUS and/or γTURC (Alfaro-Aco et al. 2020). However, neither the Aurora A kinase activating nor the importin binding part of TPX2 (Alfaro-Aco et al. 2017, Safari et al. 2021) are strictly required for Ran-GTP dependent microtubule nucleation in *Xenopus* egg extract. Moreover, TPX2 is entirely dispensable for branched microtubule nucleation in *Drosophila* cells (Verma and Maresca 2019, Goshima 2011) and recent *in vitro* reconstitutions of branched microtubule nucleation with purified human (Zhang et al. 2022) and *Drosophila* proteins (Tariq et al. 2020) were performed in the absence of TPX2. This leaves the question unanswered of how the Ran-GTP pathway controls branching microtubule nucleation mediated via the HAUS complex.

The HAUS complex is evolutionarily conserved in vertebrates, insects, plants and fungi (Edzuka et al. 2014). The hetero-octameric nature of the complex and its importance for spindle assembly was first described for *Drosophila* cells (Goshima et al. 2008). Knockdown or depletion of augmin/HAUS in cells leads to a dramatic reduction in spindle microtubule density, including in kinetochore fibers, and results in defects in spindle polarity and chromosome segregation both in *Drosophila* as well as in vertebrates and plants (David et al. 2019, Goshima et al. 2008, Lawo et al. 2009, Ho et al. 2011).

The eight subunits of the augmin complex have been reported to form two biochemically distinct subcomplexes, called tetramer T-II (containing HAUS2, 6, 7, and 8) and tetramer T-III (containing HAUS1, 3, 4, and 5) (Lawo et al. 2009). Recent cryo-electron microscopy and single particle analysis studies showed that both tetramers together form an interconnected, highly flexible Y-shaped octameric complex (with a V-shaped head and a filamentous tail) (Gabel et al. 2022, Zupa et al. 2022, Travis et al. 2022). The primary microtubule binding site has been localized to the intrinsically-disordered N-terminus of HAUS8 (Hsia et al. 2014, Wu et al. 2008), while a second, minor microtubule binding site was recently located within the HAUS6 subunit (Travis et al. 2022, Zupa et al. 2022), locating both binding sites to the V-shaped head. *In vitro* experiments suggest that several augmin complexes together recruit γTURC (Zhang et al. 2022), but how the binding of augmin to microtubules is controlled remains unknown.

Here, we report the *in vitro* reconstitution of Ran-GTP-regulated microtubule binding of the recombinant human augmin complex composed of all eight full-length subunits. We show that importins directly bind the augmin complex, thereby suppressing its microtubule binding, and that Ran-GTP relieves this inhibition of microtubule binding. We identify a previously unknown NLS motif in HAUS8 that mediates importin and microtubule binding. These findings demonstrate that the augmin complex is a direct target of the Ran-GTP pathway, suggesting that branching microtubule nucleation is directly regulated by the Ran-GTP gradient in cells.

## RESULTS

### Recombinant human HAUS complex

We expressed the human HAUS complex in insect cells using a BigBac baculovirus carrying the genes of all eight HAUS full-length subunits in codon-optimized form (Fig. 1A, S1A, B, D). HAUS2 was tagged at its C-terminus with mGFP followed by a cleavable TwinStrep-tag. The complex was purified by affinity and size exclusion chromatography, and affinity tags were removed during purification (Fig. 1B, S1C, E). Fractions eluting just after the exclusion volume of the size exclusion column were pooled (Fig. 1C) and the presence of all subunits was verified by SDS-PAGE (Fig. 1C) and western blotting (Fig. S1F), yielding a purified complex similar to a previously purified recombinant *Xenopus* and human augmin complexes (Song et al. 2018, Gabel et al. 2022). HAUS8 migrated as a double band, as in a HAUS preparation directly retrieved from cultured human cells (Zhang et al. 2022), which is due to phosphorylation, as we confirmed by mass spectrometry (Fig. S2B, C) and as observed previously (Hornbeck et al. 2015).

**Figure 1.**
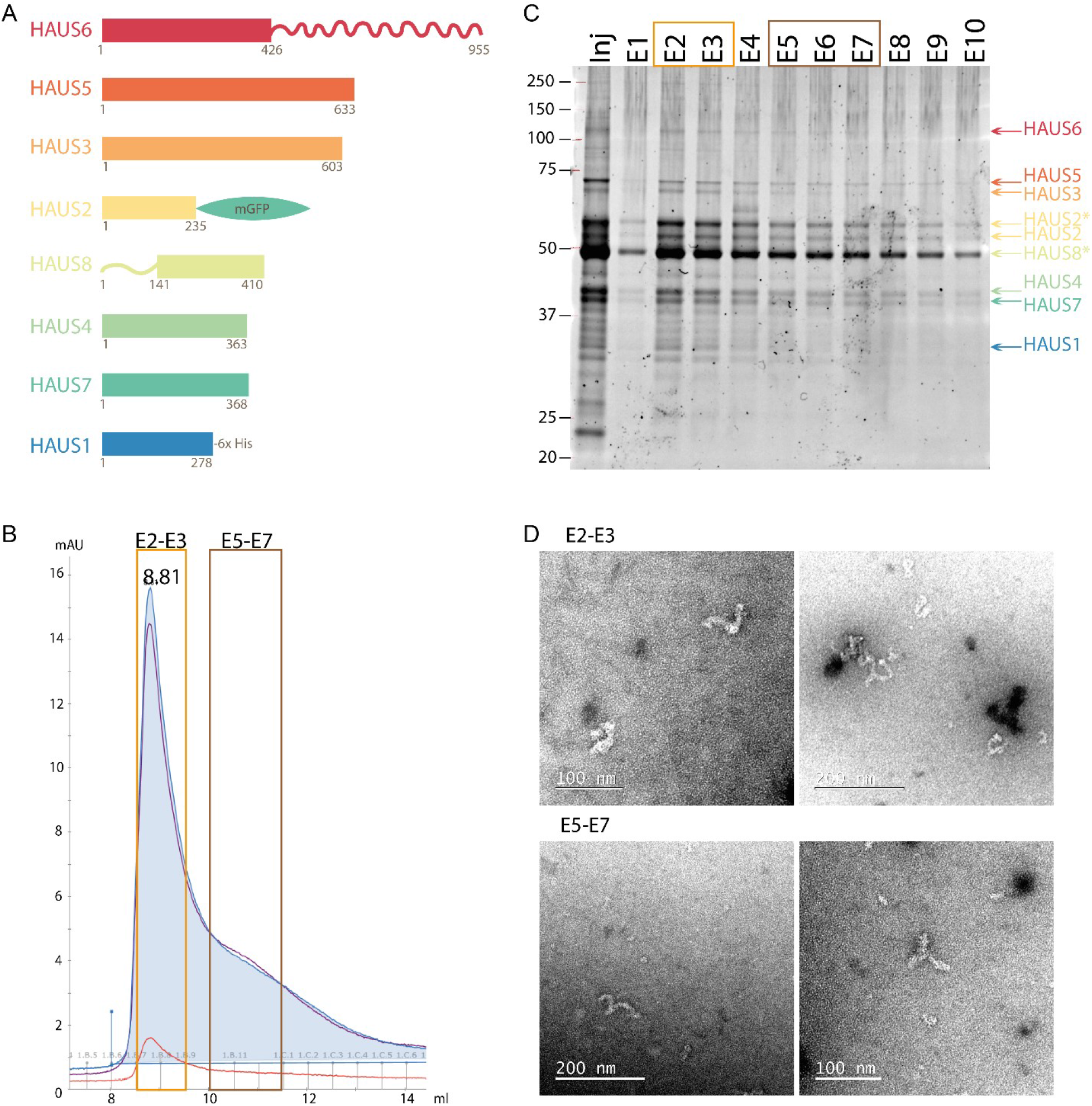
Purification and characterization of the human augmin/HAUS complex. **A.** Schematic representation of the 8 subunits of the HAUS complex. HAUS1 carries a C-terminal 3C-His6-tag, and HAUS2 has a C-terminal mGFP-TEV-TwinStrep tag. **B.** Absorbance profile of the human HAUS complex eluting from a Superose 6 Increase 10/30 size-exclusion chromatography column. The absorbance at 260, 280 and 455 nm are indicated in purple, blue and orange, respectively. The complete complex used for experiments in this study elutes in fractions E2-E3 (orange box). Fractions E5-E7 (brown box) contain mostly subcomplexes. **C.** SyproRuby stained SDS-PAGE gel of the fractions eluted from the size-exclusion chromatography column. Orange and brown boxes indicate separately pooled fractions. **D.** Negative stain electron microscopy of the different pools: fractions E2-E3 contain the complete complex, and fractions E5-E7 contain also subcomplexes.

Negative stain electron microscopy confirmed that the purified human complex had a Y-shaped structure and adopted multiple conformations (Fig. 1D top, S2A left), in agreement with recent cryo-electron microscopy structures. In our purification, the complete complex could be separated from partly assembled subcomplexes that eluted later from the size exclusion column (Fig. 1D bottom, S2A right). Together, these data indicate that the recombinant human HAUS complex consisting of all full-length subunits is correctly assembled.

### Binding of HAUS to microtubules *in vitro*

Total internal reflection fluorescence (TIRF) microscopy experiments demonstrated that the purified GFP-HAUS complex at concentrations as low as 1 nM bound all along surface-immobilized GMPCPP and taxol-stabilized TAMRA-labelled microtubules (Fig. 2A top). Increasing the GFP-HAUS complex concentration within the range of 1 nM to 20 nM increased the uniform binding to microtubules in a dose-dependent manner (Fig. 2A, B). At higher concentrations the complex started to be insoluble, preventing us from reaching saturation of binding. A similar binding efficiency was observed recently for a HAUS complex retrieved from cultured human cells (Zhang et al. 2022), whereas truncated *Xenopus* and human augmin complexes studied previously appeared to bind less efficiently, requiring using higher complex concentrations (Hsia et al. 2014, Alfaro-Aco et al. 2020).

**Figure 2.**
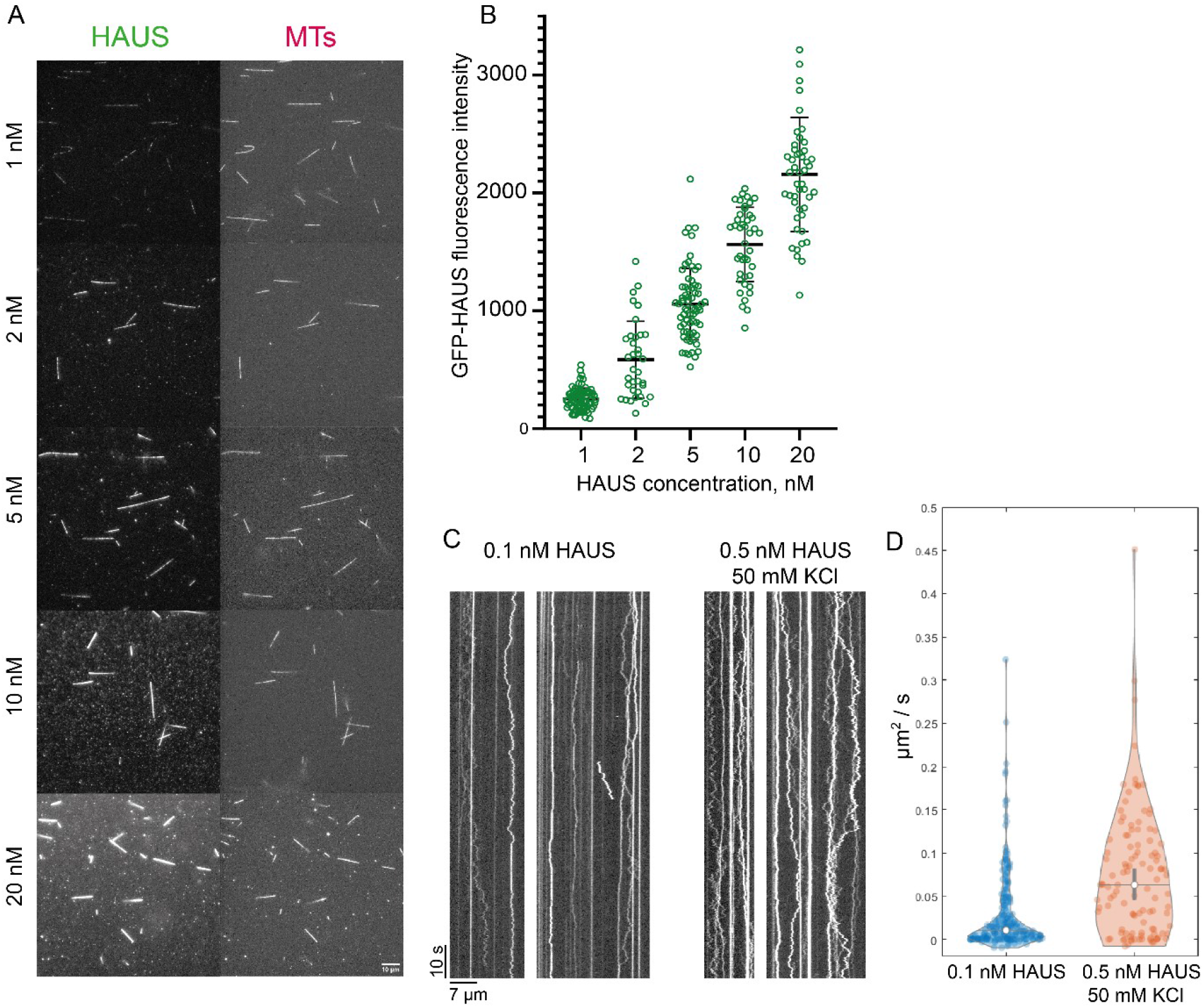
Binding of the human HAUS complex to microtubules *in vitro*. **A.** TIRF microscopy images showing binding of GFP-HAUS at the indicated concentrations (from 1 nM to 20 nM) to GMPCPP and taxol-stabilized TAMRA-microtubules (MTs). **B.** Background corrected GFP-HAUS fluorescence intensity bound to stabilized microtubules at the indicated HAUS concentrations. Black bars represent mean value ± S.D.; n = 3. **C.** TIRF microscopy kymographs showing the diffusion of single GFP-HAUS particles on microtubules. Concentrations and scale bars as indicated. **D.** Diffusion coefficients of individual tracked HAUS particles without and with added KCl. Black bars represent mean value ± S.D.; n = 3.

Next, we examined the mode of GFP-HAUS complex binding to microtubules using single-molecule TIRF microscopy imaging at lower HAUS concentrations (Fig. 2C). Kymographs (time-space plots) revealed that some complexes bound statically, whereas the majority of the complexes bound in a diffusive manner to the microtubule lattice, mostly for the entire duration of the ~ 2 min observation period. Increasing the KCl concentration by 50 mM reduced the number of observed events (Fig. S3A) which could be compensated for by increasing the HAUS concentration (Fig. 2C right). Increasing the KCL concentration also increased the measured diffusion coefficient (Fig. 2D), in agreement with an electrostatic interaction between the negatively charged microtubule surface and the positively charged unstructured N-terminal part of the HAUS8 subunit that represents the major microtubule-binding region of the complex (Wu et al. 2008). A more detailed fluorescence intensity analysis of the diffusing HAUS particles revealed the presence of a mixture of single molecules and small oligomers of up to ~ 10 molecules; larger oligomers diffused more slowly, probably due to increased friction (Fig. S3B).

### Importins bind to the HAUS complex and inhibit its microtubule binding

During the optimization of the human HAUS purification, we noted in a mass spectrometry analysis the presence of insect cell importins, suggesting that these proteins might be HAUS binding partners (Fig. S4A). We, therefore, produced purified human importin α and importin β and a constitutively active form of human Ran (RanQ69L) (Fig. S4B) to test whether they bind the HAUS complex and thereby might control its microtubule binding. The addition of importin α to a TIRF microscopy-based binding assay had no effect on microtubule binding of the HAUS complex (Fig. S4C), in agreement with importin α being self-inhibited (Lott and Cingolani 2011). However, the addition of importin β inhibited microtubule binding of HAUS in a dose-dependent manner (Fig. 3A, B, green data, Fig. S4D). A mixture of both importin α and importin β inhibited HAUS binding to microtubules even more efficiently, reaching almost complete inhibition at 100 nM importin α/β (Fig. 3C, D green data). Adding RanQ69L at a concentration of 3.5 μM to the highest tested concentrations of importin β and the mixture of both importins fully restored HAUS binding to microtubules, demonstrating that Ran-GTP can completely relieve the inhibitory effect of the importins (Fig. 3B, D orange data).

**Figure 3.**
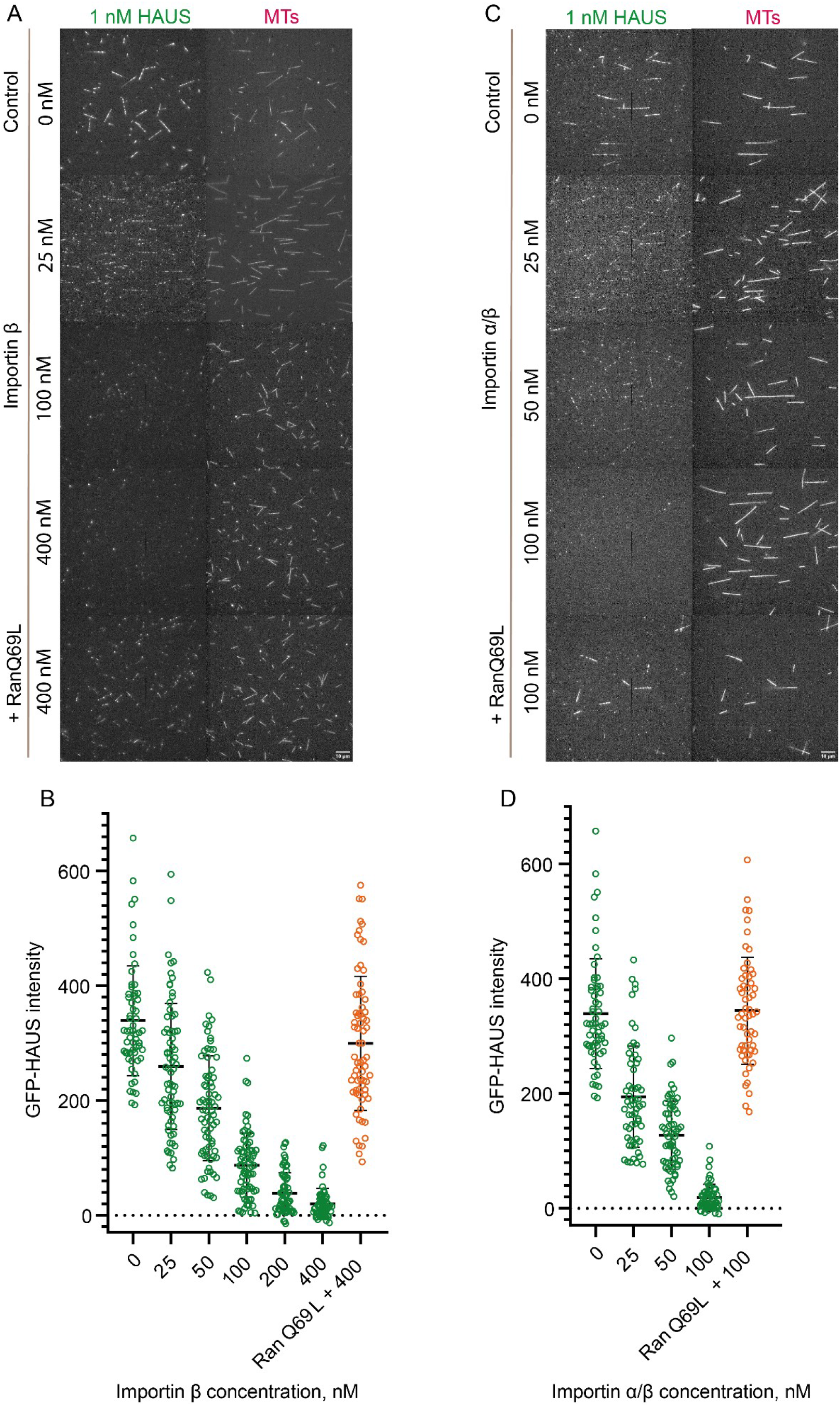
Reconstitution of the regulation of HAUS binding to microtubules by importins and Ran-GTP. **A.** TIRF microscopy images showing reduced binding of GFP-HAUS (1 nM) to TAMRA-microtubules (MTs) in the presence of increasing concentrations of importin β (25 – 400 nM). Inhibition of binding by 400 nM importin β is reverted by adding 3.5 μM constitutively active RanQ69L. **B.** Background corrected fluorescence intensities of GFP-HAUS measured on microtubules in the presence of different importin β concentrations, as indicated, and with 3.5 μM RanQ69L added to the highest importin β concentration tested (conditions as in A and Fig. S4D). Black bars represent mean values ± S.D.; n = 3. **C.** TIRF microscopy images showing reduced binding of GFP-HAUS (1 nM) to TAMRA microtubules (MTs) in the simultaneous presence of importin α and importin β (25 - 100 nM each). Inhibition of binding by 100 nM importin α/β is reverted by adding 3.5 μM RanQ69L. **D.** Background corrected fluorescence intensity of GFP-HAUS measured on microtubules in the presence of different importin α/β concentrations as indicated, and with RanQ69L added to the highest importin α/β concentration tested. Black bars represent the mean ± S.D.; n = 3.

To quantify the strength of the interaction between the HAUS complex and importin α/β, we used microscale thermophoresis and measured a dissociation constant of 13.1 ± 3.65 nM, indicating strong binding (Fig. 4A).

**Figure 4.**
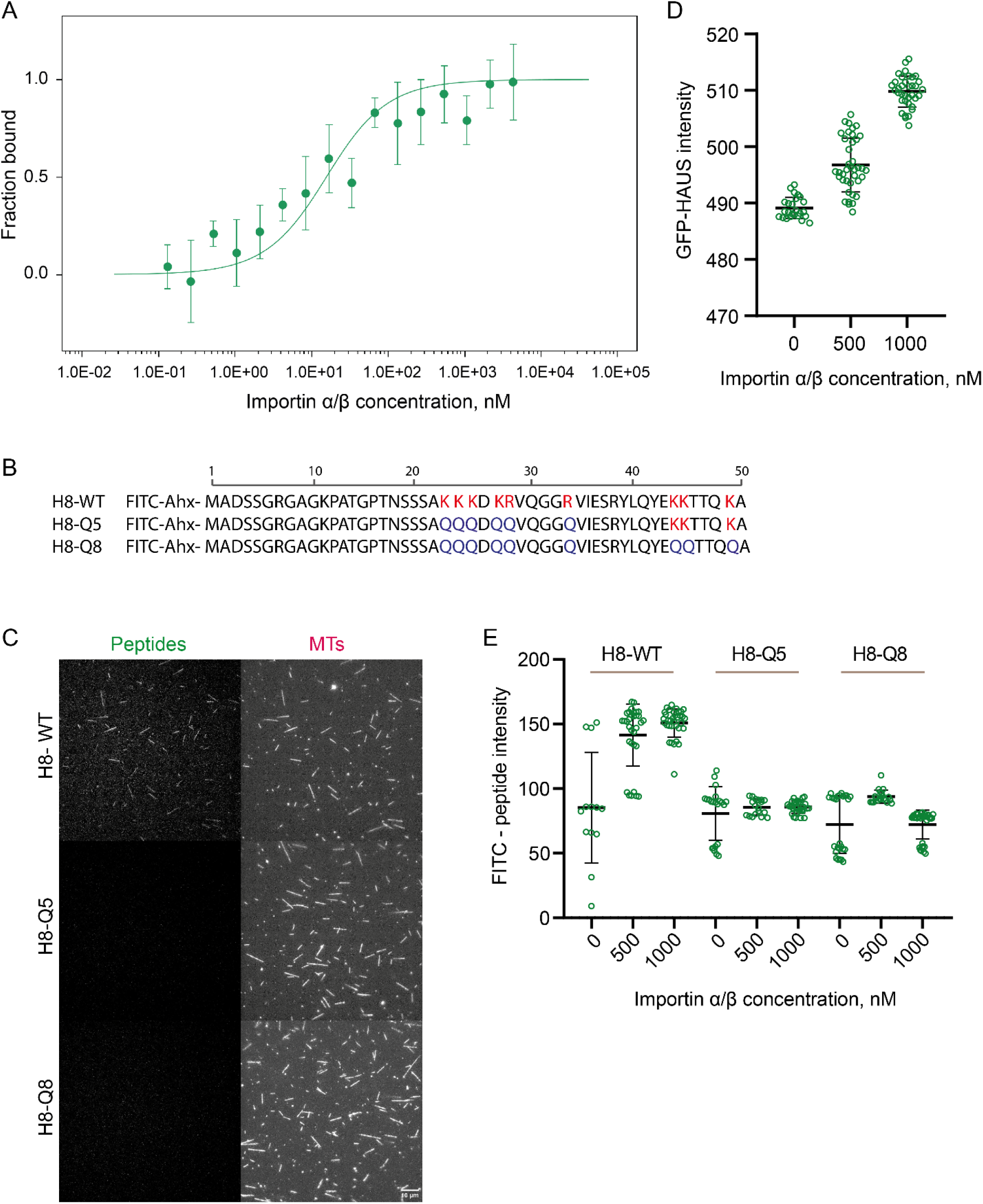
Binding of the human HAUS complex to importins and identification of the importin-regulated microtubule binding site of HAUS. **A.** Quantification of concentration-dependent binding of HAUS to importin α/β using microscale thermophoresis. GFP-HAUS (5 nM) was mixed with different concentrations of importin α/β (4.3 μM – 0.131 nM). The obtained apparent dissociation constant was 13.1 ± 3.65 nM. Data points represent the mean ± S.D.; n = 3. **B.** Sequences of synthetic peptides derived from the N-terminus of HAUS8 (amino acids 1-50). Positively charged amino acids corresponding to potential binding sites are marked in red. Amino acids mutated to glutamines are marked in blue. **C.** TIRF microscopy images showing binding of FITC-labelled wildtype peptide (100 nM) and no binding of mutated FITC-peptides (100 nM) to stabilized TAMRA-microtubules (MTs). **D.** Fluorescence intensities of GFP-HAUS complex (2.5 nM) binding to surface-immobilized importin α/β. Importin concentrations used for immobilization were as indicated. Raw fluorescence intensities are shown. Example images are shown in Fig. S5. Green data points represent intensities measured in different fields of view, black bars represent mean values ± S.D.; n = 2. **E.** Background corrected fluorescence intensities of FITC-peptides (10 nM) binding to surface-immobilized importin α/β. Importin concentrations used for immobilization were as indicated. Example images are shown in Fig. S5. Green data represent intensities measured in different fields of view, black bars represent mean values ± S.D.; n = 2.

These data demonstrate that the HAUS complex is a direct target of the Ran-GTP pathway.

### Identification of a microtubule and importin binding site in HAUS8

To identify the inhibitory importin binding site in the HAUS complex, we inspected the N-terminal unstructured part of HAUS8 that is known to contain the major microtubule binding site, although its exact location is unknown (Wu et al. 2008). Using the NLStradamus method for NLS motif prediction (Nguyen Ba et al. 2009), we identified a potential monopartite nuclear NLS motif (**KKK**D**KR**, amino acids 23 to 28 of HAUS8) that could have the ability to bind microtubules and importins in a competitive manner.

To test this hypothesis, we studied fluorescently labelled synthetic peptides that correspond to the first 50 amino acids of HAUS8 (Fig. 4B). Fluorescence microscopy showed that the peptide with the wild-type sequence (H8-WT) bound all along stabilized microtubules (Fig. 4B, 4C top), whereas a mutant in which the 5 basic amino acids of the putative monopartite NLS were changed to glutamines (H8-Q5) did not bind microtubules (Fig. 4B, C middle), like a peptide with three more mutations in a second stretch of positively charged amino acids (H8-Q8) (Fig. 4B, C, bottom). Compared to experiments with the complete HAUS complex, we used here higher concentrations of wild-type peptide for comparable binding to microtubules. These results indicate that the identified potential NLS motif forms at least part of the microtubule-binding region of HAUS8, narrowing down the part of the N-terminal extension of HAUS8 that binds to microtubules.

To investigate whether the wild-type peptide binds importins, we immobilized importin α/β on a glass surface and added increasing concentrations of the different fluorescent HAUS8 peptides, and imaged their binding by TIRF microscopy (Fig. S5). As a positive control, GFP-HAUS was used and was shown to bind to the importin surface (Fig. 4D). The wild-type peptide also bound the immobilized importins in a dose-dependent manner, in contrast to the mutated peptides that did not bind importins noticeably (Fig. 4E). These data demonstrate that the binding of importins to the identified monopartite NLS is specific. Together these data suggest that HAUS8 binding to importins and to microtubules is competitive, revealing at least part of the mechanism by which importins and Ran-GTP control the binding of the HAUS complex to microtubules.

## DISCUSSION

We showed here that the human HAUS complex is a direct target of Ran-GTP-mediated regulation. This regulation happens at the level of microtubule binding which in turn will affect the recruitment of γTuRC and possibly other nucleation stimulating factors needed to trigger chromatin-mediated branched microtubule nucleation for correct spindle assembly during cell division (Fig. 5).

**Figure 5.**
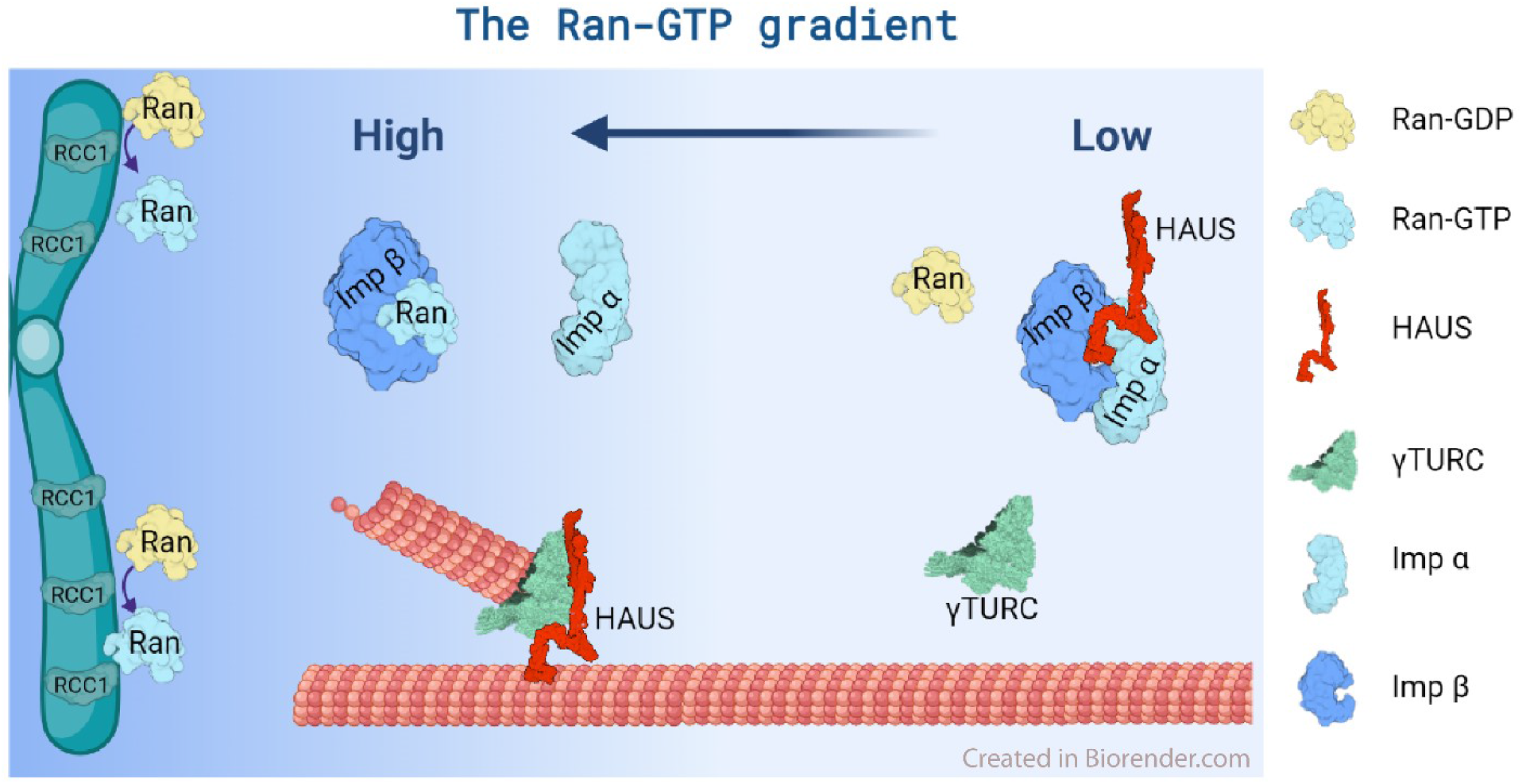
Schematic visualization of Ran-GTP-based regulation of the human HAUS complex. A Ran-GTP gradient forms around the chromosomes due to the enrichment of the exchange factor RCC1 on the chromatin. High concentrations of Ran-GTP promote the dissociation of NLS-containing protein HAUS from inhibitory importin α/β. The released HAUS can bind to microtubules and trigger branched microtubule nucleation locally in the vicinity of chromosomes.

This finding suggests an explanation why in some organisms such as *Drosophila* branched nucleation is dependent on the Ran-GTP pathway (Chen et al. 2015), but independent of TPX2 (Verma and Maresca 2019), although TPX2 is an important SAF in *Xenopus* eggs with a well-established microtubule nucleation stimulating activity (Petry et al. 2013). Our result also aligns with a recent *in vitro* reconstitution of branched microtubule nucleation using human proteins that did not include TPX2 (Zhang et al. 2022), indicating that it is not strictly required for branched microtubule nucleation in humans. Mechanistic differences may exist in different species.

We identified a previously undetected monopartite NLS motif in the N-terminal unstructured region of the HAUS8 subunit. This positively charged region is also present in HAUS8 orthologs in other species (for example in *Xenopus, Drosophila* and *Ararbidopsis* (Hotta et al. 2012)). Our observation of this NLS motif being important for microtubule binding narrows down the microtubule binding region in this part of HAUS8 (Wu et al. 2008). Recent cryo-EM studies proposed that additionally calponin homology domains identified in HAUS6 and HAUS7 might also contribute to microtubule binding of the HAUS complex (Gabel et al. 2022). Specificity of binding might be conferred by the folded calponin homology domains, and the affinity of the interaction may be controlled by the unstructured, positively charged part of HAUS8.

In agreement with our findings, a recent preprint identified two NLS motifs in the HAUS8 subunit of *Xenopus* augmin, one of which does not exist in human HAUS8 (Kraus et al. 2022). These motifs in the *Xenopus* protein were also shown to be responsible, at least in part, for importin/Ran-GTP-dependent regulation of the binding of *Xenopus* augmin to microtubules. Therefore, the basic principle of direct importin/Ran-GTP regulation of the HAUS complex appears to be conserved between human and *Xenopus*, even if details of the NLS motifs may differ, which could explain why, in contrast to the human HAUS complex, microtubule binding of the *Xenopus* HAUS complex could not be completely blocked by importins (Kraus et al. 2022).

Positively charged stretches that mediate or strengthen microtubule binding have been found in several microtubule associated proteins such as Tau (Lee et al. 1989), ASAP (Saffin et al. 2005), and NuSAP (Raemaekers et al. 2003). They are often a target of regulation, either by inhibitory importin binding (NuSAP (Ribbeck et al. 2006), HURP (Koffa et al. 2006), TPX2 (Safari et al. 2021)) or by inhibitory phosphorylation by kinases (doublecortin (Schaar et al. 2004), MAP6 (Baratier et al. 2006), Tau (Gong and Iqbal 2008)) or by other posttranslational modifications (HDAC6 (Liu et al. 2012)).

It has been reported that the N-terminal extension of human HAUS8 is also regulated by inhibitory phosphorylation by Aurora A kinase (Tsai et al. 2011). Phosphorylation of amino acids that are closely located to the NLS motif identified here, reduced microtubule nucleation in spindles of cultured human cells and impaired mitotic progression. Moreover, HAUS complex with phosphorylated HAUS8 was enriched at the spindle poles (Tsai et al. 2011). This may suggest differential regulation of augmin in different parts of the spindle controlled by the combined action of importins and Aurora A kinase.

Mass spectrometry analysis of our purified recombinant HAUS complex indicated that despite strong phosphorylation of HAUS8, its N-terminus was unphosphorylated in agreement with the observed strong HAUS complex binding to microtubules in the absence of importins. Compared to the entire HAUS complex, the HAUS8-derived peptide H8-WT (amino acids 1 −50) bound more weakly to microtubules. This is similar to the previous observation that the entire unstructured part of HAUS8 alone bound 10 times more weakly to microtubules than an augmin subcomplex containg also the calponin homology domain containing subunits HAUS 6 and HAUS7 (T-II subcomplex) (Hsia et al. 2014), further supporting the notion of several parts of the complex contributing to microtubule binding.

In agreement with previous *in vitro* reconstitutions (Zhang et al. 2022, Hsia et al. 2014), we observed diffusive microtubule binding of the human HAUS complex. Diffusion slowed down for larger HAUS oligomers. The majority of the HAUS particles were bound to microtubules for at least 2 minutes which can be compared to the measured dwell time of augmin of ~ 1 min at nucleation sites in *Drosophila* cells (Verma and Maresca 2019). Oligomerization and strong microtubule binding of the HAUS complex may be functionally important, given that more strongly bound augmin complexes observed in *Drosophila* cells were part of more active nucleation branch points (Verma and Maresca 2019). Moreover, a recent *in vitro* reconstitution of branched microtubule nucleation with human proteins also revealed HAUS oligomerization at active branch points (3 to 4 HAUS complexes per oligomer) (Zhang et al. 2022). Given the flexible structure of individual HAUS complexes, multiple complexes may have to come together to ensure the preferential geometrical orientation of HAUS/γ-TURC-nucleated microtubules with the the characteristic shallow angles observed for branched microtubule nucleation.

In conclusion, the HAUS complex is a direct target of Ran-GTP-dependent regulation which controls its binding to microtubules via importins. This sheds new light on the functioning of one of the key regulators of microtubule nucleation during cell division. Ran-GTP dependent regulation of the HAUS complex can therefore be expected to control indirectly the recruitment of γTuRC by augmin for branched microtubule nucleation. Other SAFs such as TPX2 in *Xenopus* or for example HURP in *Drosophila* may have additional roles in regulating branched microtubule nucleation by either further promoting γTuRC recruitment or its activation by other mechanisms. This will be an interesting area for further future investigation.

## MATERIALS AND METHODS

### DNA constructs

The genes of the subunits of the human augmin complex (HAUS1 (Q96CS2; NM_138443.4); HAUS2 (Q9NVX0; NM_018097.3); HAUS3 (Q68CZ6; NM_001303143.2); HAUS4 (Q9H6D7; NM_001166269.2); HAUS5 (O94927; NM_015302.2); HAUS6 (Q7Z4H7; NM_017645.5); HAUS7 (Q99871; NM_001385482.1); HAUS8 (Q9BT25; NM_033417.2)) were synthesized in codon-optimized form for expression in *Trichoplusia ni* cells, including a polyhedrin promoter and terminator overhangs of the pLIB vectors for the assembly in a baculovirus biGBac expression vector (Weissmann and Peters 2018, Weissmann et al. 2016).

The genes of HAUS1 with a C-terminal 3C-His6-tag and HAUS3 were cloned into pBiG1a; the genes of HAUS2 with a C-terminal mEGFP-TEV-TwinStrep tag, HAUS6, and HAUS7 were cloned into pBiG1b; and the HAUS4, HAUS5 and HAUS8 genes into pBiG1c. Eventually, all subunits of the HAUS complex were joined into the pBIG2abc vector by Gibson assembly (Weissmann and Peters 2018, Weissmann et al. 2016). The correct insertion of the genes was verified at each step of cloning by the restriction enzymes SwaI, PmeI, and PacI. The correct sequence of the final vector was verified by Shotgun sequencing (MGH CCIB DNA Core, MA, USA).

The bacterial expression vectors for human importin α and β and RanQ69L, all with N-terminal His6-Ztag tags, were described previously (Roostalu et al. 2015, Kronja et al. 2009).

### Protein expression and purification

#### HAUS complex

The baculovirus preparation for recombinant human HAUS complex expression was carried out according to the manufacturer’s protocol (Bac-to-Bac system, Life Technologies) using *E. coli* DH10EmBacY and Sf21 insect cells. Baculovirus-infected insect cells (BIICs) (Fig. S1D) were then frozen before cell lysis to generate stable viral stocks as described previously (Wasilko et al. 2009).

Human HAUS complex expression was induced by adding 0.8 ml of frozen BIICs per liter to a High Five insect cell culture grown to densities of ~1.2×10^6^ cells/ml. Cells were harvested 60 h post-induction by centrifugation (15 min, 1000 g, 4°C). Cell pellets were then washed with ice-cold PBS, centrifuged again (15 min, 1000 g, 4°C), frozen in liquid nitrogen, and stored at −80°C.

Pellets from 2 liters of insect cell culture were resuspended in ice-cold lysis buffer (50 mM phosphate buffer pH 7.5, 400 mM NaCl, 4 mM MgCl_2_, 5% glycerol, 2 mM TCEP, 1 mM PMSF) supplemented with protease inhibitors (Roche), DNase I (10 μg ml/ml, Sigma), PhosSTOP phosphatase inhibitor cocktail tablet (Roche) using twice the volume of buffer compared to the cell pellet. Resuspended cells were lysed by the Avestin EmulsiFlex-C5 (2 rounds). The lysate was then clarified by centrifugation (30 min, 50 000 g, 4°C), and filtered through the 1.2 μm pore filters. Avidin was added at 50 μg/ml to the lysate. The supernatant was passed through a 1 ml StrepTrap HP column (GE Healthcare), the column was washed with 20 column volumes of the lysis buffer and the protein was eluted with 3 mM desthiobiotin (Sigma, D1411).

The eluate was concentrated using an Amicon Ultra-4 concentrator (Merck Millipore, 10 kDa cut-off), and incubated overnight at 4°C with TEV and 3C proteases to remove the C-terminal StrepTagII and C-terminal 6xHis tag from the HAUS2 and HAUS1 subunits respectively. The solution was then passed over a Superose 6 Increase 10/30 size-exclusion chromatography column (GE Healthcare Life Sciences) that was preequilibrated in 50 mM phosphate buffer pH 7.5, 250 mM NaCl, 4 mM MgCl2, 3% glycerol, 0.5 mM TCEP. The collected fractions were analyzed using SDS PAGE and SyproRuby staining, and the HAUS complex containing fractions were pooled, concentrated, flash-frozen and stored in liquid nitrogen until further use. The protein concentration was measured at 280 nm using a NanoDrop One Microvolume UV-Vis spectrophotometer (Thermo Fisher Scientific).

#### Importins α and β

Importins were expressed and purified as described previously using a Protino Ni-TED resin for affinity purification, followed by removal of the N-terminal His6-Ztag by TEV protease cleavage and size exclusion chromatography using a Superdex 200 16/60 column (GE Healthcare) (Roostalu et al. 2015). The aliquoted purified protein was stored in liquid nitrogen.

#### Ran Q69L

The constitutively active Ran mutant was expressed in BL21-pLysS (Novagen) *E. coli* cells. Transformed cells were grown to a density of OD_600_ = 0.5 and induced with 0.5 mM IPTG. After overnight expression at 18°C, the cells were harvested and snap-frozen. The cells were lysed using an Avestin Emulsiflex in Ran buffer (25 mM MOPS pH 7.2, 150 mM NaCl, 1 mM MgCl_2_, 1 mM 2-mercaptoethanol), and centrifuged (30 min, 250 000 g). The supernatant was loaded on a CoCl_2_-charged 5 ml HP chelating column (GE Healthcare), eluted with Ran buffer supplemented with 300 mM imidazole, and dialyzed into Ran buffer. The Z-tag was cleaved off using His-tagged TEV protease and the solution was passed through the HP chelating column to remove the protease and uncleaved RanQ69L. 10 mM EDTA and 1 mM GTP were added and the solution was incubated on ice to allow for the loading of RanQ69L with GTP. Afterward, MgCl2 was gradually added to 10 mM and the protein was dialyzed overnight into Ran buffer. Afterward, the GTP concentration was adjusted to 1 mM and the protein was concentrated (Vivaspin). Finally, sucrose and DTT were added to final concentrations of 125 mM and 1 mM, respectively. RanQ69L was snap frozen and stored in liquid nitrogen.

#### Tubulin

Porcine tubulin was purified from the pig brain as described previously (Hyman et al. 1991, Castoldi and Popov 2003). The final buffer was BRB80 (80 mM PIPES pH 6.8, 1 mM EGTA, 1 mM MgCl_2_). The final concentration was measured at 280 nm using NanoDrop One Microvolume UV-Vis spectrophotometer.

#### Tubulin labelling

Tubulin was labelled with (5(6)-TAMRA Succinimidyl Ester (Invitrogen, C1171) for fluorescence microscopy assays according to published methods (Consolati et al. 2022).

### SDS-PAGE, western blotting

Protein samples were resolved by SDS-PAGE (NUPAGE, 10% polyacrylamide gel, Thermo Scientific) (180 V, 70 min). For SyproRuby staining, the SYPRO^®^ Ruby gel stain (ThermoFisher Scientific) was used following the manufacturer’s protocol. The stained gel was imaged using Biorad Molecular Imager Gel Doc XR+.

For immunoblotting, the gel was transferred to an iBlot PVDF membrane using the iBlot™ 2 Gel Transfer Device (ThermoFisher Scientific) under standard conditions. The membrane was blocked with 5% skimmed milk dissolved in PBS. Anti-HAUS tubulin antibodies (HAUS1 - HPA040652, Atlas antibodies, x500 dilution; HAUS2 - GTX118734, x500 dilution, GeneTex; HAUS3 - NBP3-05007, Novusbio x1000 dilution; HAUS4 - LS-C155274, Lsbio, x1000 dilution; HAUS5 - A305-827A-M, Thermofisher scientific, x1000 dilution; HAUS6 - GTX118732, GeneTex, x1000 dilution; HAUS7 - A305-557A, Thermofisher scientific, x500 dilution; HAUS8 - LS-B10617-50, Lsbio, x1000 dilution) and secondary polyclonal swine anti-rabbit antibody conjugated to HRP (P0399, Agilent Technologies, x3000 dilution) were used for the detection of HAUS subunits. The chemiluminescent signal was detected on the iBright™ FL1500 imager (ThermoFisher Scientific).

### Negative-stain electron microscopy

8 μL HAUS complex at 7 nM was added onto a transmission electron microscopy (TEM) grid and blotted after 1 minute. Then, 8 uL of uranyl acetate (2%) was deposited on the grid for 1 min before draining it off with a Whatman filter paper. TEM examination was performed with a JEM-400 (JEOL USA, Pleasanton, CA, USA) transmission electron at 120 KV. The microscope was equipped with a Gatan Inc.: Orius SC1000 CCD Camera charge-coupled device camera (Gatan, Pleasanton, CA).

### TIRF microscopy assay

Flow chambers were prepared as described previously (Skultetyova et al. 2017, Ustinova et al. 2020, Braun et al. 2011). Briefly, microscope chambers were built from silanized coverslips (Menzel 1.5H; Marienfeld High Precision 1.5H) prepared following a previously described protocol using HCl for activation (Wedler et al. 2022, Hyman 1991). Parafilm was used to space the glass coverslips to form channels of ~ 0.1 mm thickness, ~ 3 mm width, and 18 mm length. Double-stabilized (GMPCPP/taxol) microtubules prepared as described previously (Skultetyova et al. 2017, Ustinova et al. 2020, Braun et al. 2011) were attached to the glass surface in each chamber via anti-β-tubulin antibodies (Sigma-Aldrich, T7816, 20 μg/ml in PBS). Subsequently, various concentrations of mGFP-HAUS complex with or without importins and RanQ69L, at concentrations as indicated in the main text and figure legends, were added to the chamber in the binding buffer: BRB80 supplemented with 1 mM TCEP (AMRESCO, Solon, OH), 0.02 mg/ml κ-casein (Sigma, C0406), 40 mM D-glucose (Sigma, G7528), 3 mM ATP (Merck, A2383), 2 mM GTP (Jena Bioscience, NU-1047), 250 μg/ml glucose oxidase (Serva, 22778), and 60 μg/ml catalase (Merck, C-40).

### Total internal reflection fluorescence (TIRF) microscope set-up

TIRF microscopy was performed using an automated Nikon Eclipse T*i* with Perfect Focus System (Nikon, Japan), a 100x oil immersion TIRF objective (NA=1.49) (CFI SR Apo) (Nikon, Japan), and an Andor iXon 888 Ultra EMCCD camera (Andor Technology, Belfast, UK) controlled by MetaMorph software (Molecular Devices). The sample was excited using 360° TIRF illumination. The following filter combinations were used: a 488nm TIRF filter set (TRF49904; Chroma, Bellows Falls, VT) with an additional ET525/50 (Chroma) bandpass filter, and a 561nm TIRF filter set (TRF49909; Chroma) with additional ET607/70 (Chroma) bandpass filter. Visualization was done by sequential dual-color imaging (switching between 488 nm and 561 nm excitation lasers to detect the mGFP-HAUS complex and TAMRA–labeled microtubules, respectively). The image acquisition rate for the observation of HAUS single-molecule movements was 1 frame per 113 ms. The microscope chamber was kept at 30°C using OkoLab temperature control (OkoLab).

### TIRF microscopy image analysis

Experimental data were processed with Fiji (Schindelin et al. 2012, Rueden et al. 2017) and Fiesta (Ruhnow et al. 2011).

To measure overall fluorescence intensities of microtubule-bound mGFP-HAUS, we followed a previously described procedure (Ustinova et al. 2020). Briefly, areas around microtubules were manually determined; the fluorescence intensity of all pixels in these areas were then measured in the mGFP-HAUS channel. After background subtraction the average fluorescence intensity per pixel was calculated for each microtubule area, corresponding to one experimental data point. The data were plotted using GraphPad Prism software. Mean values and standard deviation were calculated for each experimental condition.

Overall fluorescence intensities of mGFP-HAUS or FITC-peptides to surface-immobilized importins were measured as described above, but measuring the intensities in the entire field of view.

For diffusion measurements, mGFP-HAUS particles were tracked using FIESTA. All tracks were manually verified and projected on an averaged path using a custom-written MATLAB script (MathWorks, Natick, MA)(MATLAB 2018). From these positions, the one-dimensional diffusion coefficient for each complex was estimated using a covariance-based estimator (Vestergaard et al. 2014). The intensity of each complex was estimated using the averaged integrated intensity of the Gaussian fit (using only the first 20 frames of each track). The arbitrary intensity count was converted to the number of fluorescent molecules by dividing the intensity counts by the average integrated intensity of single mGFP-HAUS molecules (estimated in a separate photobleaching experiment).

### Microscale thermophoresis

A two-fold dilution series of importins α/β was prepared in BRB80, 1 mM TCEP, 0.02 mg/ml κ-casein, 3 mM ATP, and 2 mM GTP. An equal amount of 10 nM mGFP-HAUS complex in the same buffer was added to the importin α/β dilution series (final importin concentrations were 4.3 μM–0.131 nM). The reactions were incubated for 5 min at room temperature. The Monolith NT.115 microscale thermophoresis instrument with premium coated capillaries (NanoTemper, Munchen, Germany) was used to measure binding curves. The excitation power was set to 40% and the MST power was set to 60%. The results were processed with MO Affinity Analysis.

### Peptide synthesis

A 50-residue peptide sequence from the N-terminal part of HAUS8 and 2 mutated versions were synthesized:

wildtype: MADSSGRGAGKPATGPTNSSSAKKKDKRVQGGRVIESRYLQYEKKTTQKA,
mutant 1: MADSSGRGAGKPATGPTNSSSAQQQDQQVQGGQVIESRYLQYEKKTTQKA,
mutant 2: MADSSGRGAGKPATGPTNSSSAQQQDQQVQGGQVIESQYLQYEQQTTQQA.

The peptides were assembled in a Liberty Blue instrument running 0.1 mmol-scale solid phase synthesis Fmoc protocols on Rink amide ProTide resin. Side chain protections were NG-2,2,4,6,7-pentamethyldihydrobenzofuran-5-sulfonyl (Arg), t-butyl (Ser, Thr, Asp, Glu), t-butyloxycarbonyl (Lys, Trp) and trityl (Asn, Gln). Double couplings with 5-fold molar amounts of Fmoc-amino acid, Oxyma and diisopropylcarbodiimide in DMF, with microwave activation at 90°C, were used at every cycle. Fluorescent labelling at the N-terminus was done by manual coupling of a 5(6)-carboxyfluorescein unit. Further details on the synthesis, cleavage, purification and analytical documentation of similar peptides can be found in Carrera-Aubesart et al., 2022(Carrera-Aubesart et al. 2022).

### Mass spectrometry

Gels bands were destained (100 mM NH_4_HCO_3_ in 40% acetonitrile), reduced (DTT), alkylated (iodoacetamide), and dehydrated for trypsin digestion (Promega, V5113). Obtained peptide mix was desalted with a MicroSpin C18 column (The Nest Group, Inc) prior to LC-MS/MS analysis. Samples were analyzed using an Orbitrap Fusion Lumos or Orbitrap Eclipse mass spectrometer (Thermo Fisher Scientific, San Jose, CA, USA) coupled to an EASY-nLC 1200 (Thermo Fisher Scientific (Proxeon), Odense, Denmark). Peptides were separated by reversed-phase chromatography using a C18-column (Thermo Scientific, San Jose, CA, USA).

Digested bovine serum albumin (New England Biolabs, P8108S) was analyzed between each sample to avoid sample carryover and to assure the stability of the instrument. QCloud (Chiva et al. 2018) has been used to control instrument longitudinal performance during the project.

Acquired spectra were analyzed using the Proteome Discoverer software suite (v1.4 or 2.5, Thermo Fisher Scientific) and the Mascot search engine (v2.6, Matrix Science (Perkins et al. 1999)). The data were searched against a SwissProt_Human database plus *Trichoplusia ni* reference proteome (UP000322000) (35205 entries) or SwissProt_Human database plus sf21 UniProt reference proteome (UP000240619) (20821 entries) plus a list (Beer et al. 2017) of common contaminants and all the corresponding decoy entries. Oxidation of methionine and N-terminal protein acetylation were used as variable modifications whereas carbamidomethylation on cysteines was set as a fixed modification. The false discovery rate (FDR) in peptide identification was set to a maximum of 5%.

### Visualization

For the preparation of the scheme, the online platform BioRender was used. The following protein structures from PDB were used for schematic visualization - Ran (3GJ0), importin α (6IW8), importin β (1QGK), γTURC (6V6S), HAUS complex (7SQK).

## ACKNOWLEDGEMENTS

We thank Johanna Roostalu for providing purified importins and RanQ69L, Silvia Speroni and Raquel Garcia-Castellanos for technical support and advice. We thank Martí de Cabo Jaume from the Servei de Microscòpia of Universitat Autònoma de Barcelona for electron microscopy support, the Protein Technologies Facility of the Center for Genomic Regulation for microscale thermophoresis support, Guadalupe Espadas-Garcia from the CRG/UPF Proteomics Unit which is part of the Spanish National Infrastructure for Omics Technologies (ICTS OmicsTech) for performing the proteomics analysis, and David Andreu Martínez from the Peptide Synthesis Facility, Department of Medicine and Life Sciences (MELIS), Pompeu Fabra University, Barcelona for the peptide synthesis. MELIS is a member of the “María de Maeztu” Program for Units of Excellence in R&D of the Spanish Ministry of Science and Innovation.

